# The mitochondrial pyruvate carrier (MPC) complex is one of three pyruvate-supplying pathways that sustain Arabidopsis respiratory metabolism

**DOI:** 10.1101/2021.02.24.432685

**Authors:** Xuyen H. Le, Chun-Pong Lee, A. Harvey Millar

**Affiliations:** The University of Western Australia, School of Molecular Sciences; The ARC Centre of Excellence in Plant Energy Biology, 35 Stirling Highway, Crawley, Perth 6009, Australia

## Abstract

Malate oxidation by plant mitochondria enables the generation of both oxaloacetate (OAA) and pyruvate for tricarboxylic acid (TCA) cycle function, potentially eliminating the need for pyruvate transport into mitochondria in plants. Here we show that the absence of the mitochondrial pyruvate carrier 1 (MPC1) causes the co-commitment loss of its orthologs, MPC3/MPC4, and eliminates pyruvate transport into Arabidopsis mitochondria, proving it is essential for MPC complex function. While the loss of either MPC or mitochondrial pyruvate-generating NAD-malic enzyme (NAD-ME) did not cause vegetative phenotypes, the lack of both reduced plant growth and caused an increase in cellular pyruvate levels, indicating a block in respiratory metabolism, and elevated the levels of branched-chain amino acids at night, a sign of alterative substrate provision for respiration. ^13^C-pyruvate feeding of leaves lacking MPC showed metabolic homeostasis were largely maintained except for alanine and glutamate, indicating that transamination contributes to restoration of the metabolic network to an operating equilibrium by delivering pyruvate independently of MPC into the matrix. Inhibition of alanine aminotransferases (AlaAT) when MPC1 is absent resulted in extremely retarded phenotypes in Arabidopsis, suggesting all pyruvate-supplying enzymes work synergistically to support the TCA cycle for sustained plant growth.

## Introduction

Pyruvate occupies a pivotal node in the regulation of carbon metabolism as the end product of glycolysis and a major substrate for mitochondrial respiration. Mitochondrial pyruvate oxidation is catalysed by the pyruvate dehydrogenase complex (PDC) to provide carbon skeletons to the tricarboxylic acid (TCA) cycle, and its depletion leads to extensive changes in metabolism, abnormal organ development and severe growth retardation in plants (Ohbayashi et al., 2019; Yu et al., 2012). PDC and malate dehydrogenase (MDH) work cooperatively to provide substrates, acetyl-CoA and oxaloacetate (OAA) respectively, for citrate synthesis (Selinski and Scheibe, 2019). In terms of kinetics, malate production by MDH is favoured over OAA formation (Hüdig et al., 2015), therefore in order to drive the reaction towards OAA and maintain the continuity of respiration, OAA needs to be continuously removed by NAD-ME (Tronconi et al., 2010b; Tronconi et al., 2010a), citrate synthase (Schmidtmann et al., 2014) and a putative OAA exporter (Haferkamp and Schmitz-Esser, 2012). Malate oxidation by plant mitochondria enables the generation of both OAA (via MDH) and pyruvate (via NAD-ME) for TCA cycle function, and in isolated plant mitochondria malate-feeding enables a much faster respiratory rate than pyruvate alone. This indicates that pyruvate import may be slower than malate import, and based on this it has long been thought that the need for pyruvate transport across the inner mitochondrial membrane in plants may be minimal (Day and Hanson, 1977). In the following decades, this concept has been further tested by various *in vitro* biochemical approaches (Edwards et al., 1998; Jenner et al., 2001) and *in vivo* ^13^C-labelling studies (Tcherkez et al., 2005; Tcherkez et al., 2008; Tcherkez et al., 2009; Lehmann et al., 2016), but genetic evidence to prove it had been lacking.

The availability of full genome information and insertional mutants in Arabidopsis enabled significant advances in our understanding the fate of malate in plant mitochondria. Insertional loss-of-function of both mitochondrial MDH enzymes resulted in a slow growth phenotype and an elevated leaf respiration rate (Tomaz et al., 2010). Insertional loss-of-function of both mitochondrial NAD-ME subunits in Arabidopsis caused a significant diversion of excess malate to amino acid synthesis at night without affecting vegetative growth (Tronconi et al., 2008). Gene insertion-based reduction in PDC activity caused a greatly retarded vegetative phenotype in Arabidopsis (Ohbayashi et al., 2019). This suggests that mitochondrial pyruvate supply and oxidation are both important for optimal plant growth.

The transport of glycolytically-derived pyruvate into the mitochondrial matrix is carried out by the mitochondrial pyruvate carrier (MPC). In yeast, a functional MPC complex is composed of a heterodimer of a core subunit (MPC1) and a regulatory unit (MPC2, MPC3) (Tavoulari et al., 2019). Yeast without MPC1 grows slowly in medium without amino acids and has a lower pyruvate transport (Bricker et al., 2012; Herzig et al., 2012). *MPC* mutation in *Drosophila* is lethal with a sugar-only diet and caused impaired pyruvate oxidation-linked metabolism under a standard diet. Human *MPC1* harbouring a single point mutation resulted in lactic acidosis and hyperpyruvatemia due to impaired pyruvate oxidation, while knock-out of MPC1 in mice led to embryonic lethality (Brivet et al., 2003). From these reports, it can be concluded that MPC1 is an essential component for supplying pyruvate to mitochondria in yeast and mammals. In Arabidopsis, four MPC isoforms are present, with Arabidopsis MPC1 (At5g20090) showing the highest homology with yeast MPC1 (Shen et al., 2017). The role of Arabidopsis MPC in pyruvate transport has been proven indirectly through complementation of a yeast mutant (Li et al., 2014). In plants, lack of MPC has only been linked to phenotypes of cadmium sensitivity and drought tolerance (Li et al., 2014; Shen et al., 2017; He et al., 2019). Single or higher order knockouts of MPC isoforms do not cause any obvious growth defect under normal conditions (He et al.,2019), indicating that unlike in other eukaryotes, pyruvate transport is probably not essential for respiratory pyruvate supply in Arabidopsis. Apart from the direct transport of glycolysis-derived pyruvate into mitochondria and malate oxidation, pyruvate can also be supplied from alanine through the combined actions of cytosolic and mitochondrial alanine aminotransferases (AlaATs) (Miyashita et al., 2007) and a yet-to-be characterised mitochondrial alanine carrier (Passarella et al., 2003; Bender and Martinou, 2016). However, the role of AlaATs in reversibly converting pyruvate to alanine has only been reported during hypoxia and recovery from hypoxia (Ricoult et al., 2006; Miyashita and Good, 2008; Diab and Limami, 2016).

MPC in yeast and mammalian systems can only facilitate pyruvate transport from the cytosol to the mitochondrial matrix and not vice versa (Bricker et al., 2012; Herzig et al., 2012). Pyruvate export from mitochondria in cancer cells is mediated by a different class of transporter called the monocarboxylate carrier MCT1 (Hong et al., 2016). Under some circumstances in plants, the export of pyruvate from mitochondria is needed for PEP synthesis in the cytosol by pyruvate orthophosphate dikinase to respond to water stress in rice seedlings (Netting, 2002) or to support the recycling of carbon intermediates and CO_2_ fixation in Crassulacean acid metabolism (CAM) (Hartwell et al., 2016) and C_4_ plants (Rao and Dixon, 2016). However, the identity of the plant mitochondrial pyruvate exporter is currently unknown. It remains unclear which MPC isoforms facilitate pyruvate transport in plants, which direction of transport they facilitate and how pyruvate transport activity via MPC is co-regulated with other pyruvate-supplying pathway in contributing to the plant mitochondrial pyruvate pool.

In this study, we show the primary role of MPC1 in pyruvate import into Arabidopsis mitochondria using a reverse genetic approach in combination with direct evidence from *in vitro, in organello*, and *in vivo* analyses. We obtained evidence that plant mitochondrial pyruvate import is mainly mediated by MPC1, which is needed to maintain pyruvate supply for *Arabidopsis* respiratory metabolism. NAD-ME can independently supply pyruvate to the TCA cycle in *Arabidopsis* mitochondria, but our data indicate that it has a relatively small contribution to pyruvate-linked metabolism *in vivo*. Instead, the action of AlaATs provides an important functional backup for MPC complex and these enzymes most likely work in a cooperative manner to supply mitochondrial pyruvate needed for sustaining respiration at night in plants. These data provide an insight into the flexibility of pyruvate-supplying pathways in plants to efficiently use and extract energy from the cellular carbon repertoire in different environments.

## Results

### MPC1 is essential for mitochondrial pyruvate carrier complex accumulation in Arabidopsis and its absence causes the co-commitment loss of MPC3 and MPC4

In Arabidopsis, *MPC1*, *MPC3* and *MPC4* are ubiquitously expressed in all tissue types while *MPC2* is expressed exclusively in flowers (Waese et al., 2017). MPC2, MPC3 and MPC4 share about 60%-80% amino acid sequence identity, whereas MPC1 is the least similar to these isoforms (<33%) but shares the highest sequence identity with yeast MPC1 (Schwacke et al., 2003; Li et al., 2014). Thus, MPC1 is the most promising candidate as a non-redundant core component of the plant MPC complex. A T-DNA insertion line, *mpc1,* was previously confirmed to be a homozygous mutant that contained an insertion 400bp upstream of the start codon (Shen et al., 2017; Supplementary Figure 1A). *mpc1* does not exhibit an obvious vegetative phenotype under long day conditions (Supplementary Figure 1B). Quantitative RT-PCR analysis of whole leaf and quantitation of selected peptides from isolated mitochondria by selective reaction monitoring - mass spectrometry (LC-SRM-MS) revealed that *mpc1* is unable to produce full length *MCP1* transcript or accumulate MPC1 protein, confirming that it is a MPC1 knockout mutant (Figure 1). Even though no obvious change in *MPC3* and *MPC4* transcript abundance was observed in *mpc1*, MPC3 and MPC4 proteins were completely absent in isolated mitochondria (Figure 1B). It is possible that *MPC3* and *MPC4* are transcribed but their protein products may be unstable, leading to their degradation. When a MPC1 transgene was expressed under the control of its native promoter in *mpc1* plants (*mpc1/gMPC1*), both MPC3 and MPC4 abundances were restored in mitochondria (Figure 1A). Thus, MPC1 is required for the assembly of the MPC complex.

**Figure 1.**
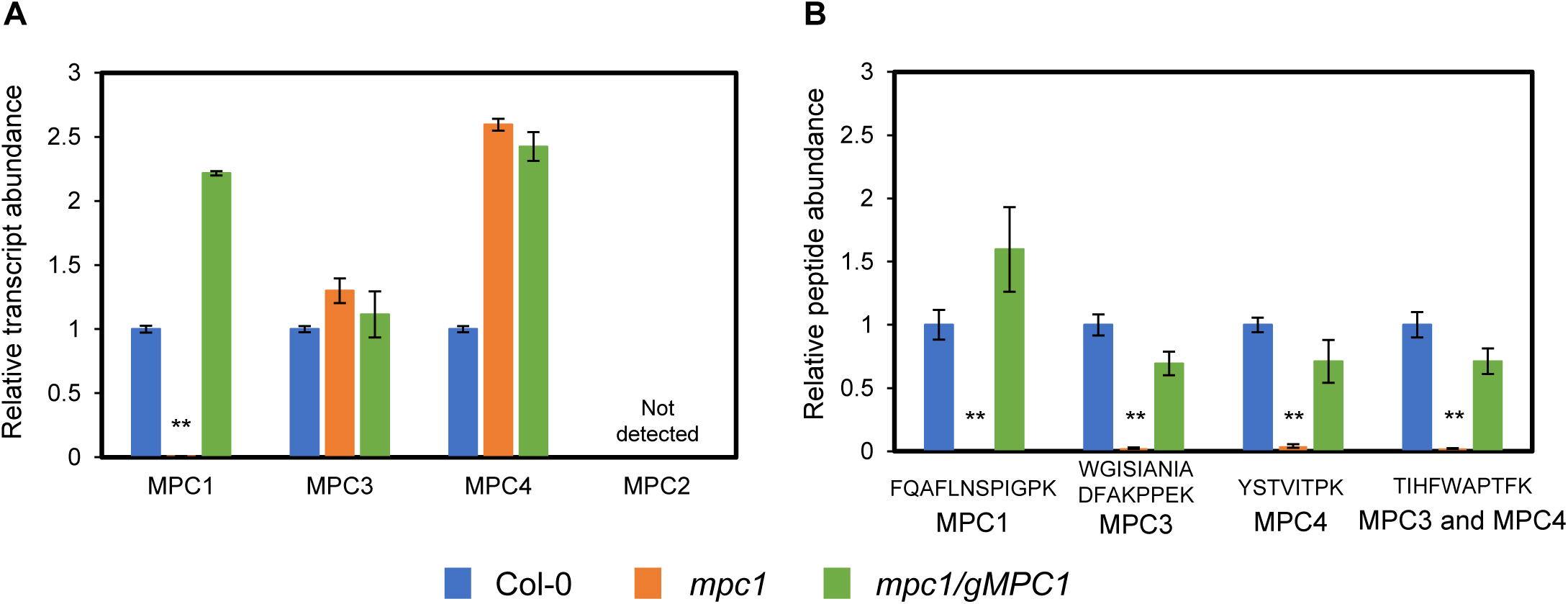
MPC1 is required for the assembly of mitochondrial pyruvate carrier complex in Arabidopsis. (A) Relative transcript abundance of *MPC*s in Col-0, *mpc1* and *mpc1/gMPC1*. Transcript abundance was measured using qPCR of *MPC1, MPC2, MPC3, MPC4* inter-exon sequences. Primers are provided in Supplementary Table 1. *ELF1α* was used as a housekeeping control. (B) Relative protein abundance of MPC isoforms in Col-0*, mpc1* and *mpc1/gMPC1*. Protein abundances were determined by quantifying peak area of unique peptides using LC-SRM-MS. FQAFLNSPIGPK was unique for MPC1, WGISIANIADFAKPPEK was unique for MPC3, YSTVITPK was unique for MPC4. The abundance of a mitochondrial VDAC1 peptide was used as a housekeeping control. Each data point represents an averaged value from three biological replicates with error bars indicating standard error. Significant differences between *mpc1*, Col-0 and *mpc1/gMPC1* are denoted by asterisks based on Student’s t-tests (**, p < 0.001).

### MPC1 is required for pyruvate-dependent respiration and is a target of the pyruvate transport inhibitor UK-5099

Evidence that plant MPC1 functions as a mitochondrial pyruvate importer to date is based on yeast complementation analysis (Li et al., 2014) and sequence conservation with yeast and mammal homologues. To provide direct experimental proof for its role in respiration and pyruvate transport, we used isolated mitochondria as an *in vitro* model system to monitor MPC function by manipulating respiratory substrate supply. We first measured the ability of isolated mitochondria to oxidise pyruvate in the presence of ADP and cofactors. Pyruvate as the sole substrate was unable to drive significant oxygen consumption, probably due to the lack of OAA replenishment. Respiratory assays at pH 7.2 have been shown previously to minimize malic enzyme conversion of exogenous malate to pyruvate in isolated mitochondria (Willeford and Wedding, 1987). When malate alone was supplied at pH 7.2 in our experiments, only 20% of the oxygen consumption rate in the presence of malate and saturating pyruvate was measured (27 ± 1.6 nmol/min/mg protein, compared to 149 ± 23 nmol/min/mg). Titrations with different non-saturating concentrations of pyruvate revealed a ∼70% reduction in oxygen consumption rate (called pyruvate-dependent OCR) by *mpc1* mitochondria compared to wildtype (Figure 2), suggesting that *mpc1* has a diminished ability to oxidise pyruvate for mitochondrial respiration. This could be the result of (i) altered PDC abundance and/or activity, (ii) defects in TCA cycle and/or electron transport chain (ETC) functions and/or (iii) inability to import pyruvate. We found that the relative abundance and activity of PDC and the relative abundance of TCA cycle enzymes showed no difference between Col-0 and *mpc1* (Supplementary Figure 2A, 2C). In addition, the relative abundance of ETC components was unaffected by the loss of MPC1 (Supplementary Figure 2B), thus the possibility of a defect in mitochondrial function other than pyruvate import was unlikely.

**Figure 2.**
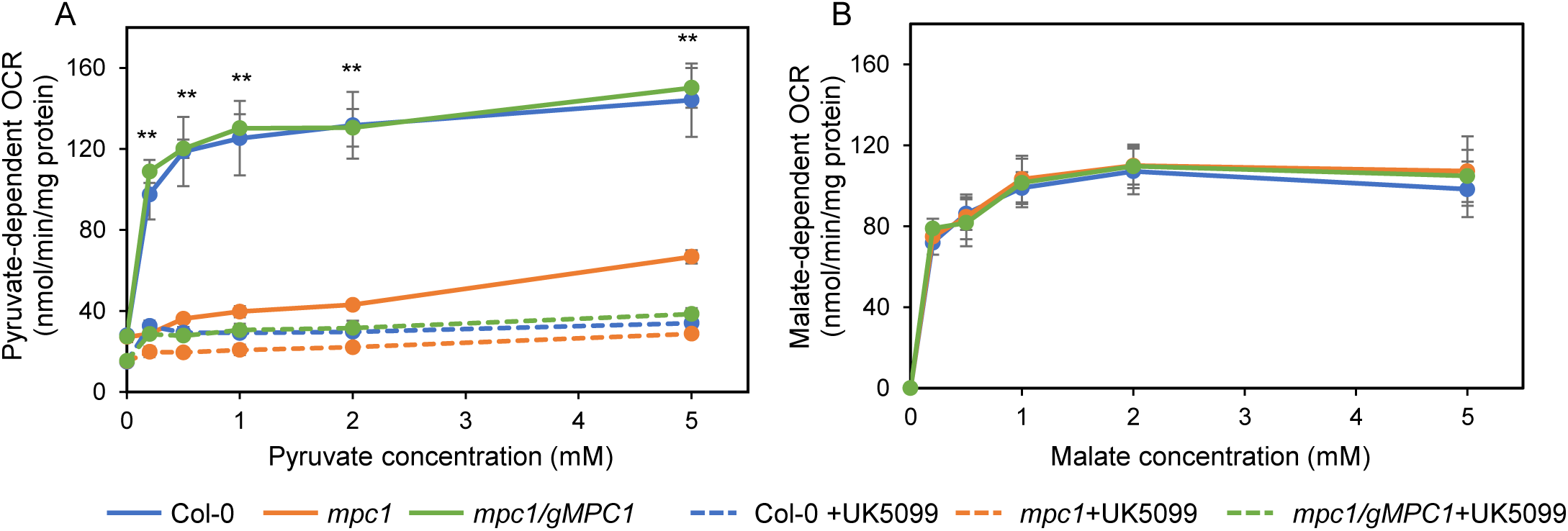
MPC1 is required for pyruvate-dependent respiration of Arabidopsis mitochondria. Mitochondrial respiration in the presence of ADP measured as oxygen consumption rate (OCR) was assayed at varying concentrations (A) of pyruvate at pH 7.2 supplemented with 0.5 mM malate or (B) of malate at pH 6.4. UK5099 was used as a pyruvate carrier inhibitor where shown. Each data point represents an averaged value from four biological replicates with error bars indicating standard error. Significant differences between *mpc1*, Col-0 and *mpc1/gMPC1* are denoted by double asterisks based on Student’s t-tests (**, p < 0.001).

As an additional control, we measured malate-dependent respiration at pH 6.4 to promote NAD-ME activity and found no difference between wildtype, mutant and complemented line (Figure 2B).

When we measured mitochondrial pyruvate-dependent respiration in the presence of UK-5099, a known inhibitor for MPC in mammals (Halestrap, 1975), we found *mpc1* mitochondria were insensitive the UK-5099 treatment (Figure 2A). In contrast, mitochondria isolated from wildtype and *mpc1/gMPC1* were sensitive to UK-5099 treatment, showing pyruvate dependence of oxygen consumption diminished to a rate similar to untreated *mpc1* mitochondria. This confirmed that the presence of MPC1 was required for UK-5099-sensitive pyruvate-dependent respiration in Arabidopsis mitochondria.

### MPC1 is required for pyruvate import into plant mitochondria

To directly demonstrate whether pyruvate uptake is defective in *mpc1* mitochondria, we provided isolated mitochondria with both unlabelled malate and ^13^C_3_-pyruvate in the presence of ADP and other cofactors (see Methods) at pH 7.2, followed by SRM-MS analysis to monitor the depletion of these metabolites in the extra-mitochondrial space (Figure 3). Figure 3A, 3B and 3F provides a detailed explanation of the experimental setup and the ^13^C incorporation patterns from the provided substrates into TCA cycle intermediates. The incorporation of ^13^C_3_-pyruvate into TCA cycle intermediates by freshly isolated mitochondria was tracked by measuring the level of metabolites in the extra-mitochondrial space (Figure 3B-E). The amount ^13^C_3_-pyruvate decreased linearly in the extra-mitochondrial space of wildtype and *mpc1/gMPC1* mitochondria, while pyruvate uptake rate was negligible in *mpc1* mitochondria, resulting in a significant difference in net pyruvate uptake (Figure 3C and Supplementary Figure 3A). This showed that the absence of MPC1 effectively abolished pyruvate import into the mitochondrial matrix.

**Figure 3.**
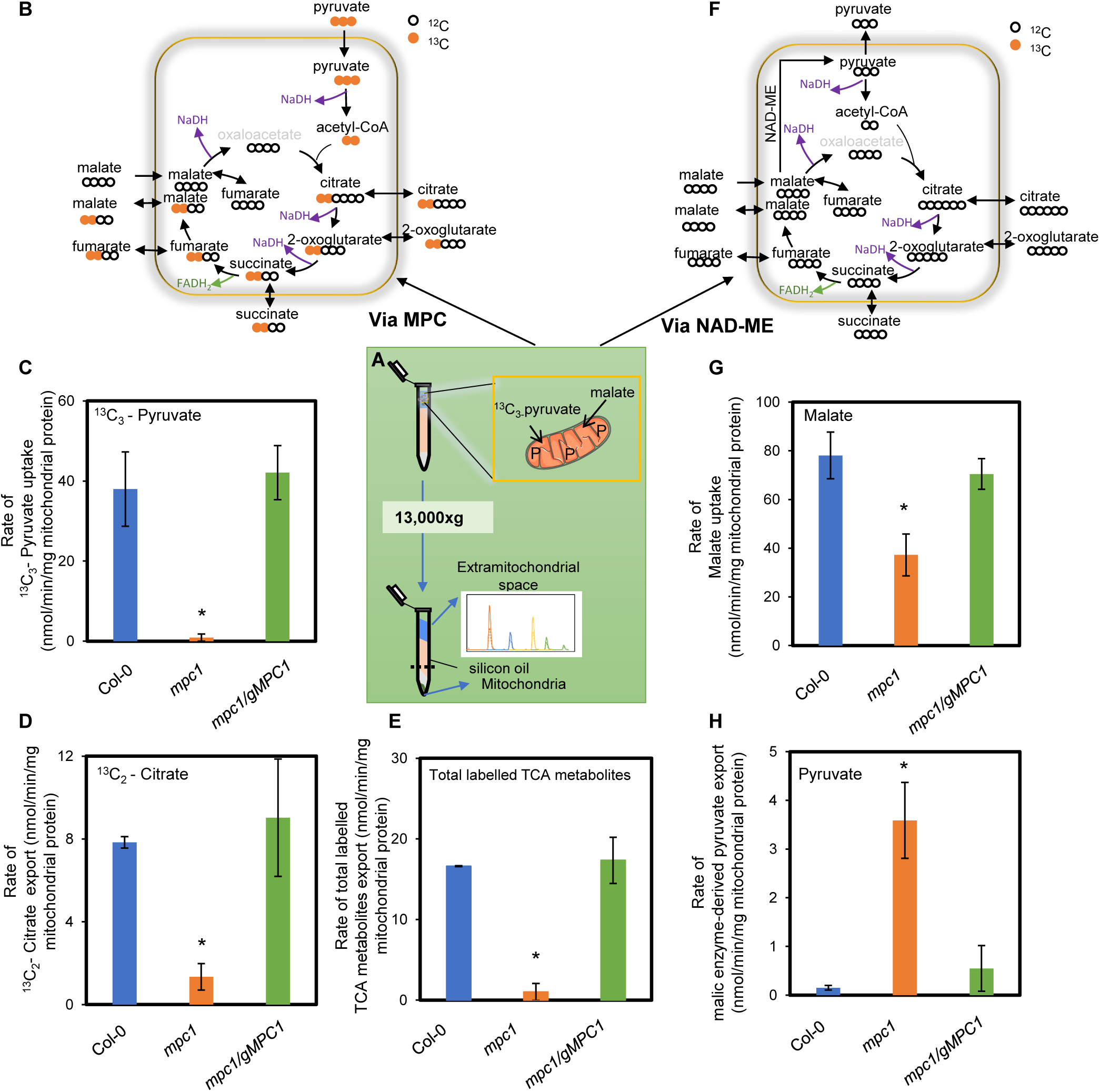
MPC1 is required for the uptake and usage of pyruvate by *Arabidopsis* mitochondria. Experimental design of combined malate and ^13^C_3_-pyruvate feeding of isolated mitochondria at pH 7.2 (A) showing the isotopic incorporation patterns of labelled pyruvate into TCA cycle intermediates along with the steps that generate NADH for consumption by the ETC via MPC (B) and NAD-ME (F). Movements of organic acids across mitochondrial membranes are also shown. Bar graphs show the (C) uptake rate of ^13^C_3_-pyruvate, (D) export rate of labelled citrate, (E) export rate of total labelled TCA metabolites made from imported pyruvate (G) uptake rate of unlabelled malate, and (H) rate of unlabelled pyruvate made via NAD-ME. Quantification was carried out using SRM-MS to directly assess substrate consumption and product generation of substrate-fed mitochondria after separating mitochondria from the extra-mitochondrial space by centrifugation through a single silicon oil layer. The rates were calculated from time course values of metabolite concentration recorded in the extra-mitochondrial space after varying incubation periods. Each data point represents averaged value from three or more replicates with error bars indicate standard error. Significant differences between *mpc1*, Col-0 and *mpc1/gMPC1* are denoted by asterisks based on Student’s t-tests (*, p < 0.05).

Mitochondria fed with substrates and cofactors to drive full TCA cycle operation have been shown to generate TCA intermediates in excess which are rapidly exported to the extramitochondrial medium (Brailsford et al., 1986). We have recently demonstrated that it is possible to assess the function of mitochondrial carriers in isolated mitochondria by following such conversion rates of supplied respiratory substrate(s) using SRM-MS analysis (Lee et al., 2020). ^13^C_2_-citrate accumulated linearly in the extra-mitochondrial space of wildtype and *mpc1/gMPC1* mitochondria while in *mpc1* only 15% of that rate was recorded (Figure 3D).

Treatment of mitochondria from all three genotypes with UK-5099 resulted in a drastic reduction in labelled citrate production, similar to that observed in *mpc1* (Supplementary Figure 3B). The total labelled carbon incorporated into measured TCA cycle intermediates from ^13^C_3_-pyruvate was significantly higher in wildtype compared to *mpc1* (Figure 3E), which confirmed *mpc1* could not sustain the same rate of supply of pyruvate in the matrix for TCA cycle-driven respiration (Figure 2). These results could be reproduced in a label swap experiment using unlabelled pyruvate and ^13^C_4_-malate as substrates, generating completely independent isotopic patterns but the same conclusions with regards to the rate of pyruvate transport in *mpc1* (Supplementary Figure 4 and 5). Taken together, these data support the claim that MPC1 is an essential subunit of the MPC complex in Arabidopsis responsible for pyruvate uptake into the matrix of plant mitochondria.

### Internal pyruvate synthesis by mitochondrial NAD-dependent malic enzyme is a compensation pathway for the loss of MPC1

In the same combined malate and ^13^C_3_-pyruvate feeding of isolated mitochondria experiment, we were also able to trace the amount of malate that was imported into the mitochondria and how it was consumed (Figure 3F-H). The amount of malate taken up by *mpc1* mitochondria was found to be half of that by wildtype and *mpc1/gMPC1* mitochondria (Figure 3G). This could be due to a lack of pyruvate in *mpc1* mitochondria to remove OAA, which could then sequentially result in a feedback inhibition of MDH, an increased malate concentration in the matrix and a reduced demand for malate uptake.

Unlike yeast and mammals where MPC1 loss caused a range of growth defects (Bricker et al., 2012; Herzig et al., 2012), the absence of MPC1 did not result in growth delay in *Arabidopsis* (Shen et al., 2017). It is therefore very likely that alternative pyruvate-supplying pathways exist in plants, with at least one of them being more active in plants than in yeast or mammals. Apart from feeding carbon through OAA via malate dehydrogenase, malate can also be converted into pyruvate to contribute to the matrix pyruvate pool (Figure 3F). The mitochondrial NAD-ME pathway is the most likely candidate given its close association with the TCA cycle, its use of malate as primary substrate, its ability to provide pyruvate in the matrix without a transport step, and evidence that its loss results in a readjustment in TCA cycle flux at night (Tronconi et al., 2008). Malate oxidation by NAD-ME in intact isolated mitochondria can generate a significant amount of pyruvate (Day, 1980) which can be assayed in the extra-mitochondrial medium. We found the rate at which pyruvate was synthesised from malate and exported to the extra-mitochondrial space was significantly higher in *mpc1* compared to wildtype and *mpc1/gMPC1* (Figure 3H), even though they have similar maximal NAD-ME activities (Supplementary Figure 2D). Similar results were observed using unlabelled pyruvate and ^13^C_4_-malate as substrates in the label swap experiment (Supplementary Figure 4 and 5). These data indicate that NAD-ME substantively provided an alternative internal pyruvate source when MPC1 was absent.

### The absence of MPC in NAD-ME knock-out plants results in a vegetative phenotype due to limited pyruvate supply as a respiratory substrate

The elevation of NAD-ME-derived pyruvate in *mpc1* mitochondria suggested the role of MPC1 could be enhanced in NAD-ME knock-out plants. To investigate this, we generated a triple knock-out by crossing *mpc1* and *nad.me1.1 x nad.me2.1* (*me1.me2)* plants to assess their combined effect on growth and metabolic phenotype *in planta*. Plants homozygous for all three mutations *nad.me1.1 x nad.me2.1 x mpc1* (*me1.me2.mpc1*; Supplementary Figure 2A) displayed significantly decreased vegetative growth rate compared to wildtype and other genotypes (Figure 4, Supplementary Figure 6). *me1.me2.mpc1* plants were noticeably smaller at the 4-leaves stage onwards even though the number of leaves were identical. At later developmental stages (Stage 1.10, 5.10 and 6.00), *me1.me2.mpc1* plants were about two to six days delayed. The size of mutant leaves did not reach the full diameter of Col-0 after one week of bolting. The retarded phenotype was observed in both long day and short-day growth conditions (Supplementary Figure 6C), however it could be restored to a wildtype-like phenotype upon re-introduction of the MPC1 transgene under the control of its native promoter (*me1.me2.mpc1/gMPC1*; Supplementary Figure 6D). These results showed that MPC is an important component for plant growth and development when NAD-ME activity is impaired.

**Figure 4.**
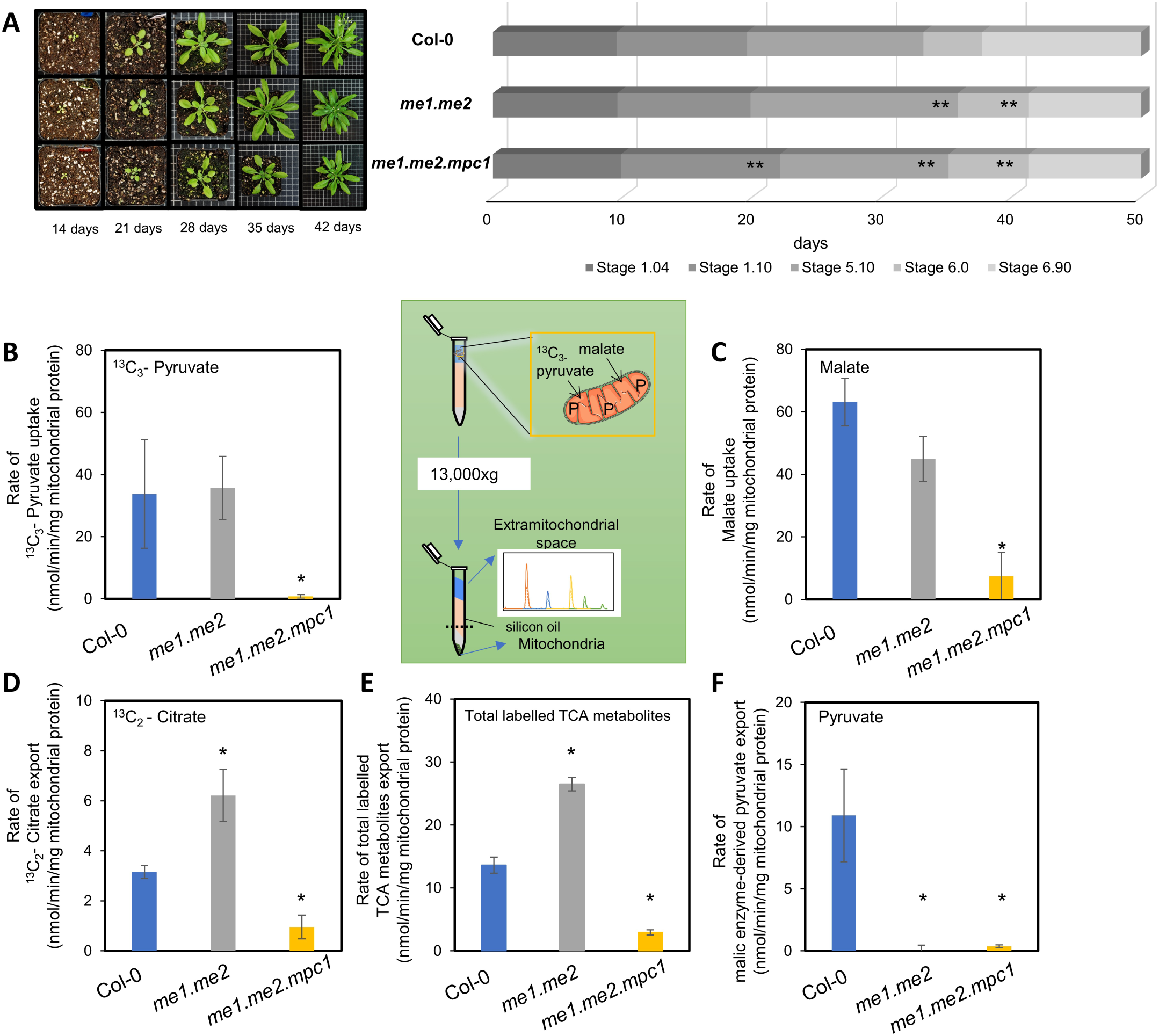
Vegetative and respiratory phenotypes of NAD-ME knock-out and MPC x NAD-ME knockouts. (A) Quantitative phenotyping of *me1.me2* and *me1.me2 x mpc1* plants in a soil-based experiment (n = 15) in long day conditions (16h light/8h dark, 60% humidity). Photos were taken at 14 days, 21 days, 28 days, 35 days and 42 days after sowing to track plant growth. Representative top views of various genotype rosettes are shown. Each gradient bar represents distinct growth stages: 1.04 - four rosette leaves > 1 mm; 1.10 - ten rosette leaves > 1 mm; 5.10 - first flower bud visible; 6.00 - first flower opened; 6.90 - flowering complete. Significant differences between tested genotypes and Col-0 are indicated based on Student t-tests and denoted by asterisks (**, p < 0.01). Bar graphs show the uptake rate of ^13^C_3-_pyruvate (B); uptake rate of malate (C); export rate of citrate (D) and export rate of total labelled TCA metabolites synthesised via MPC (E) and export rate of pyruvate synthesised via NAD-ME (F) (including ^13^C_2_-citrate, ^13^C_2_-2-oxoglutarate, ^13^C_2_-succinate and ^13^C_2_-malate). Mitochondria were incubated in 500 µM ^13^C_3_-pyruvate and 500 µM unlabelled malate in the presence of ADP at pH 6.4. Metabolic reaction was stopped by centrifugation through a single silicon oil layer in which the mitochondrial pellet was separated from the extra-mitochondrial medium. The rates were calculated from time courses of metabolites concentration recorded in the extra-mitochondrial space that were quantified using LC-SRM-MS as outlined in Methods. Each data point represents an averaged value from three or more replicates with error bars indicating standard error. Significant differences between mutants and Col-0 are indicated based on Student’s t-tests and denoted by asterisks (*, p < 0.05).

The functional importance of MPC is also highlighted by how mitochondria isolated from NAD-ME double knock-out plants consumed externally-fed malate and pyruvate. To show this, isolated mitochondria were fed with both ^13^C_3_-pyruvate and malate at pH 6.4 to maximise the rate of NAD-ME activity. At pH 6.4, MPC activity of Col-0 appeared to be similar to that at pH 7.2 (Figure 3C and Figure 4B). *me1.me2* plants did not have a vegetative phenotype (Figure 4A) and therefore were expected to have wildtype pyruvate transport activity using MPC. Indeed, malate and ^13^C_3_-pyruvate were taken up by wildtype and *me1.me2* mitochondria at a similar rate (Figure 4B and 4C). However, while wildtype mitochondria converted imported malate to pyruvate via NAD-ME, no rate of pyruvate export could be measured in *me1.me2*. Instead, *me1.me2* mitochondria showed an even higher consumption rate of ^13^C_3_-pyruvate than wildtype, as evidenced by an increased rate of ^13^C_2_-citrate synthesis and export (Figure 4D). This enhanced conversion into downstream labelled TCA intermediates, such as succinate and malate, was consistent with *me1.me2* having a rate of imported pyruvate consumption higher than wildtype (Figure 4E, Supplementary Figure 6). In contrast, the loss of both MPC and NAD-ME in *me1.me2.mpc1* resulted in the inability of isolated mitochondria to import and oxidise externally-fed pyruvate and malate, presumably due to the accumulation of OAA in the matrix. Mitochondria from these triple knockout plants displayed nearly no U-^13^C-pyruvate transport and relatively low malate transport, 0.75 ± 0.58 and 7.4 ± 7.6 nmol/min/mg respectively, compared to 33.7 ± 17.4 and 63.1 ± 7.6 nmol/min/mg protein in wildtype mitochondria (Figure 4B). Reduced incorporation of the supplied substrates into *me1.me2.mpc1* mitochondria was confirmed by the limited rate of conversion into subsequent TCA intermediates (Supplementary Figure 7).

### Alanine is a third source of mitochondrial pyruvate in *Arabidopsis*

Compared to the substantial growth phenotype caused by a decrease in PDC activity (Ohbayashi et al., 2019), the phenotype of *me1.me2.mpc1* plants is comparatively mild. This suggests that there could be a third source of mitochondrial pyruvate to support the development of mutant plants lacking both NAD-ME and MPC. To explore which metabolite could be responsible, we measured pool sizes of primary metabolites over a light/dark cycle. Metabolomics revealed that *mpc1* leaves accumulated 4-6 fold higher levels of alanine throughout the diurnal cycle compared to wildtype (Figure 5). An increase in the abundances of valine, leucine and leucine were also observed in *mpc1*. These amino acids are most likely synthesised as a consequence of glycolytic pyruvate that cannot enter into the mitochondrial matrix (Singh and Shaner, 1995). Pyruvate and malate were slightly increased in abundance in *mpc1* at night but no overall abundance change in TCA cycle intermediate was observed (Supplementary Figure 8). In contrast, knockout of NAD-ME led to elevated pyruvate and 2-oxoglutarate (2-OG) levels at night. *me1.me2.mpc1* appeared to exhibit the combined effect of *mpc1* and *me1.me2* on metabolite abundances (Figure 5A). In addition, knockout of both MPC1 and NAD-ME magnified the pyruvate accumulation to 4-fold compared to wildtype, which was consistent with a block in pyruvate-dependent mitochondrial metabolism (Figure 5A).

**Figure 5.**
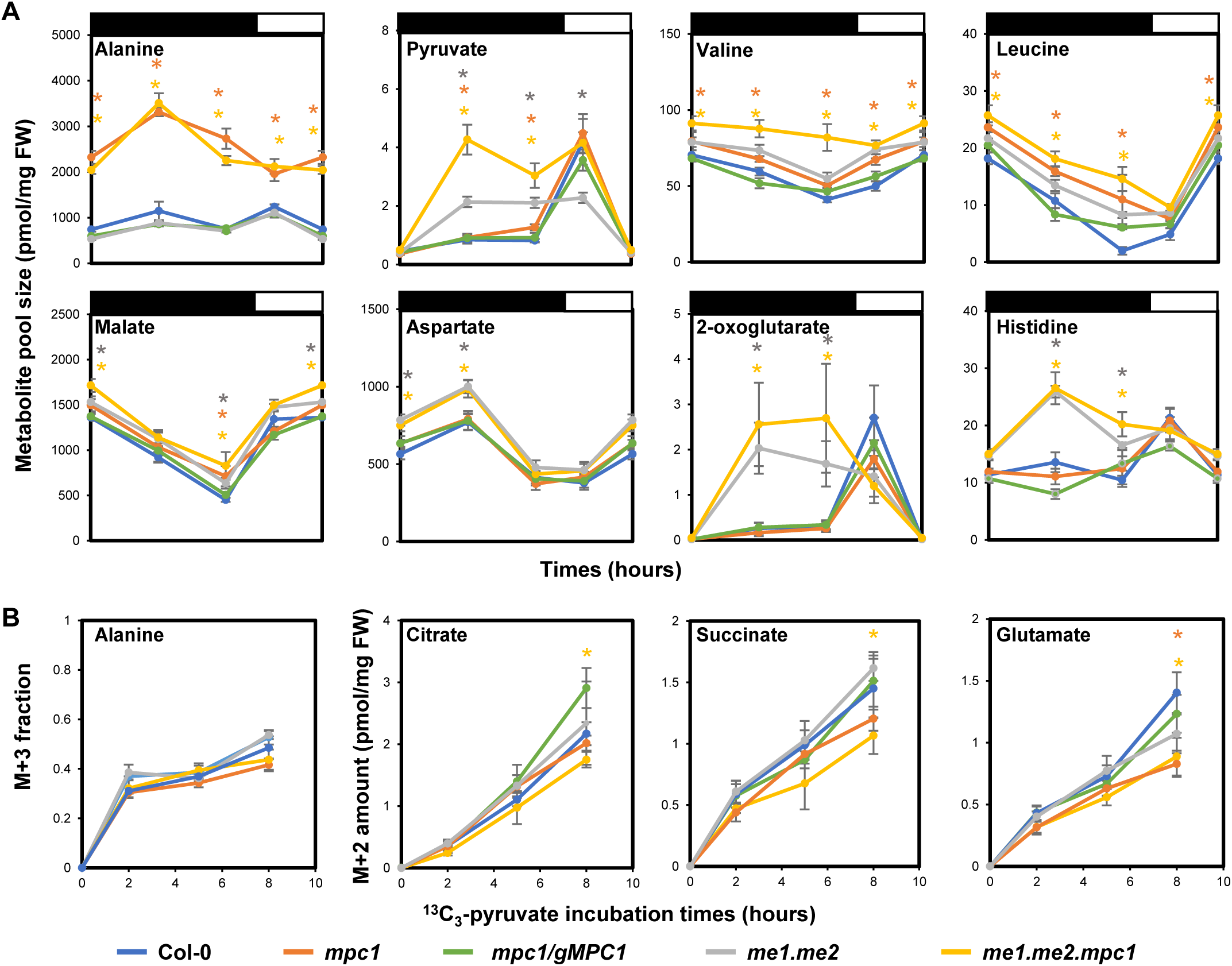
Alanine accumulation and usage helps supply pyruvate in the absence of MPC and NAD-ME. (A) Diurnal changes of representative metabolites of Col-0 (blue), *mpc1* (orange), *mpc1/gMPC1* (green), *me1.me2* (grey), *me1.me2.mpc1* (yellow). Leaf discs were collected from 6 weeks old plants grown under short day conditions. Samples were collected at 1; 8; 15 hours after the switch to darkness and 4 hours after the switch to light. The length of dark and light periods are indicated by black and white bars, respectively. The unit of metabolite absolute amount (vertical axis) is pmol/mg fresh weight. Each data point represents an averaged value from at least six replicates with error bars indicating the standard error. Significant differences between tested genotypes vs Col-0 are indicated based on Student’s t-tests and denoted by corresponding coloured asterisks. (B) Time courses of ^13^C-labelling into metabolites using ^13^C_3_-pyruvate in darkened leaf discs. Leaf discs were collected from 6 weeks old plants grown under short day conditions and incubated in 20 mM ^13^C_3_-pyruvate solution for 2; 5 and 8 hours, respectively. Incorporation patterns of ^13^C_3_-pyruvate into TCA cycle intermediates via PDC are similar to information shown in Figure 3A. [M+3] alanine fraction and absolute abundances of [M+2] citrate, [M+2] succinate and [M+2] glutamate are shown. Means ± S.E (n=4). Significant differences between mutants and Col-0 are indicated based on Student’s t-tests and denoted by corresponding coloured asterisks (*, p < 0.05).

To probe the rate of pyruvate incorporation into primary metabolism in intact tissue, we fed ^13^C_3_-pyruvate to leaf discs in the dark. Amongst five genotypes, only *me1.me2.mpc1* showed a slightly lower rate of ^13^C-incorporation into citrate and succinate which could be attributed to the deficiency in mitochondrial pyruvate utilisation due to the lack of NAD-ME and MPC activities (Figure 5B). The M+3 alanine fraction (directly produced from amination of ^13^C_3_-pyruvate) showed no significant change in relative distribution (Figure 5B) while the M+3 amount increased more significantly in *mpc1* and *me1.me2.mpc1* than that in wildtype (Supplementary Figure 8), which indicates that the increased alanine pool size was due to a higher demand for this amino acid rather than blockage in alanine-requiring enzymatic reactions. Similar results were also observed following ^13^C_6_-glucose labelling of leaf discs (Supplementary Figure 9). These data provide evidence that alanine-2-OG transamination is a third pathway for providing pyruvate to sustain mitochondrial TCA cycle metabolism and respiration. Mitochondrial AlaAT abundance measured at wildtype level in *mpc1* and *me1.me2.mpc1* (Supplementary Figure 2) suggests that the increase in alanine use was not driven by enzyme concentration. Indeed, the significantly lower amount of labelled glutamate (Figure 5B) and the elevated level of 2-OG pool size at night (Figure 5A) in *me1.me2.mpc1* plants indicate an altered equilibrium in total cellular transamination activities compared to the wildtype. Such a change in 2-OG:glutamate in plants lacking MPC1 could be a consequence of increased demand for alanine transamination reactions in mitochondria and cytosol. This pathway via AlaAT appeared to carry sufficient pyruvate flux for plant metabolism as label incorporation into most TCA cycle metabolites was maintained at wildtype levels, except for citrate and succinate (Figure 5B, Supplementary Figure 8).

### Plants lacking MPC are hypersensitive to inhibition of alanine aminotransferases by cycloserine

The existence of a yet-to-be identified mitochondrial alanine carrier (Passarella et al., 2003) could enable alanine to be imported from the cytosol into mitochondria. As AlaATs in mitochondria and the cytosol can inter-convert pyruvate and alanine, it would be possible to bypass MPC by replacing pyruvate with alanine as a primary entry point for glycolytic products to enter mitochondrial metabolism. Using the established AlaAT inhibitor cycloserine (Wong et al., 1973; Cornell et al., 1984; Duff et al., 2012), we subjected *mpc1*, *me1.me2* and *me1.me2.mpc1* and wildtype plants to chemical inhibition of AlaATs to assess the contribution of different pyruvate sources to seedling development (Figure 6). The *me1.me2.mpc1* plants displayed only a moderate change in rosette size under normal conditions, suggesting that AlaAT alone can largely support seedling growth. In the presence of 0.5µM cycloserine, however, the rosette size and root length of *me1.me2.mpc1* and *mpc1* was significantly reduced compared to that of wildtype plants. These phenotypes could be complemented by reintroducing MPC1 transgene into the *mpc1*-carrying mutants (Figure 6). *me1.me2.mpc1* was more sensitive than *mpc1* to this inhibitor, as shown by the significant difference in their rosette size and root length. This could be interpreted as evidence that NAD.ME activity in *mpc1* accounts for a small percentage of pyruvate generation *in vivo*. By contrast, *me1.me2* seedlings were insensitive to cycloserine treatment, indicating that MPC alone was sufficient to supply the bulk of pyruvate needed for sustaining seedling growth.

**Figure 6.**
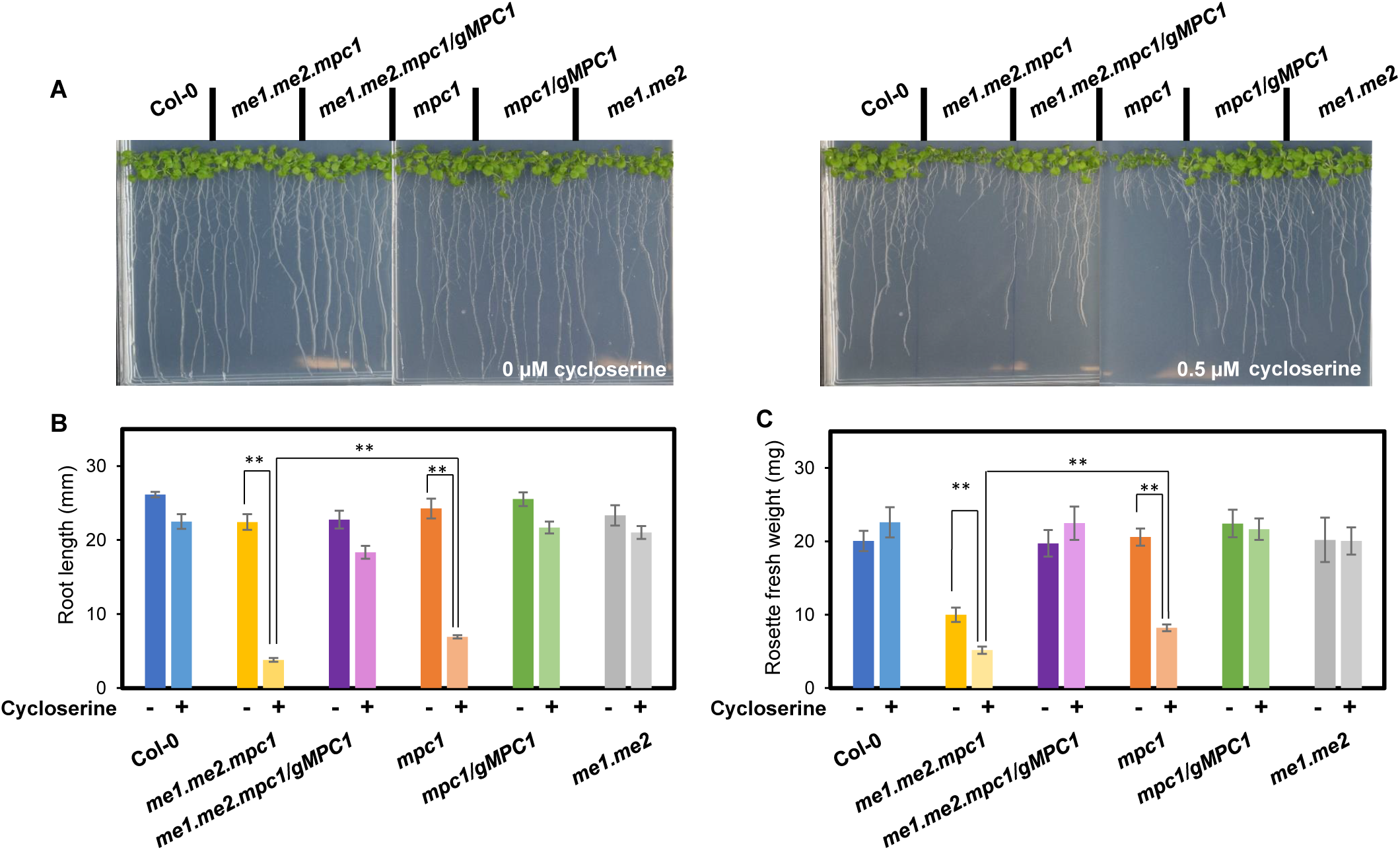
Phenotypes of MPC1 and/or NAD.ME mutants in response to cycloserine treatment. Seeds were sterilised and placed on ½-MS plates in the absence (left) or presence (right) of 0.5µM cycloserine and grown under long day conditions (16h light/8h dark, 60% humidity) (A). Representative photos are taken after 2 weeks. Bar graphs showing root length (B) and rosette fresh weight (C) of the six genotypes measured after 2 weeks. Each data point represents an averaged value from ten replicates with error bars indicating the standard error. Significant differences between genotypes are indicated based on Student’s t-tests and denoted by asterisks (**, p < 0.01).

Taken together, our data show that MPC and NAD-ME can both independently supply pyruvate to the TCA cycle in Arabidopsis mitochondria. However, when both these pyruvate sources are absent, metabolite pool sizes are still largely maintained (Supplementary Figure 7) with the exception of alanine utilisation, and a mild change in plant development is observed. Blocking AlaATs severely affects growth in *me1.me2.mpc1* and *mpc1* while it has little effect on wildtype plants. This suggests pyruvate-alanine cycling via cytosolic and mitochondrial AlaATs is another important pathway that works cooperatively with MPC and NAD-ME to maintain mitochondrial pyruvate supply for dark respiration in plants.

## Discussion

Mitochondrial pyruvate transport has not been considered as the dominant pathway for supplying oxidative substrates to respiration in plants for decades (Day and Hanson, 1977; Tcherkez et al., 2005; Araújo et al., 2012; Lehmann et al., 2016) since malate alone can provide pyruvate and OAA concomitantly within the matrix to drive the TCA cycle operation (Macrae and Moorhouse, 1970, Selinski and Scheibe, 2019). Although MPC1 has been recently shown to impact cadmium tolerance and stomatal opening in plants (Shen et al., 2017; He et al., 2019), no studies have been conducted in plants to prove that the MPC complex transports pyruvate into mitochondria or demonstrate its roles in mitochondrial respiratory metabolism. In this study, we provided the first genetic and biochemical evidence for MPC1 being the core carrier isoform responsible for the assembly and function of mitochondrial pyruvate import complex in plants. In yeast, the presence of yeast MPC1 stabilizes yeast MPC2 and MPC3 through physical interaction as functional heterodimers (Tavoulari et al., 2019). In a similar manner, the loss of MPC1 in *Arabidopsis* resulted in the absence of both MPC3 and MPC4 (Figure 1B). This explains why *mpc1* and higher-order mutants carrying *mpc1* allele produce the same cadmium-dependent short-root phenotypes as *mpc1* (He et al., 2019). Our results show the MPC acts only as a mitochondrial pyruvate importer, as mitochondrial matrix generated pyruvate could still be readily exported to the extra-mitochondrial space by *mpc1* mitochondria (Figure 3G, Supplementary Figure 3 and 4). This suggests the existence of other mitochondrial pyruvate transporter(s) that export pyruvate from the mitochondrial matrix which is consistent with evidence that carriers other than MPC are responsible for mitochondrial pyruvate export in mammals (Hong et al., 2016; Chinopoulos, 2020). More crucially, our study also provides strong evidence that mitochondrial NAD-ME and cytosolic and mitochondrial AlaATs operate cooperatively with MPC to maintain the mitochondrial pyruvate pool *in vivo* in Arabidopsis (Figure 7).

**Figure 7.**
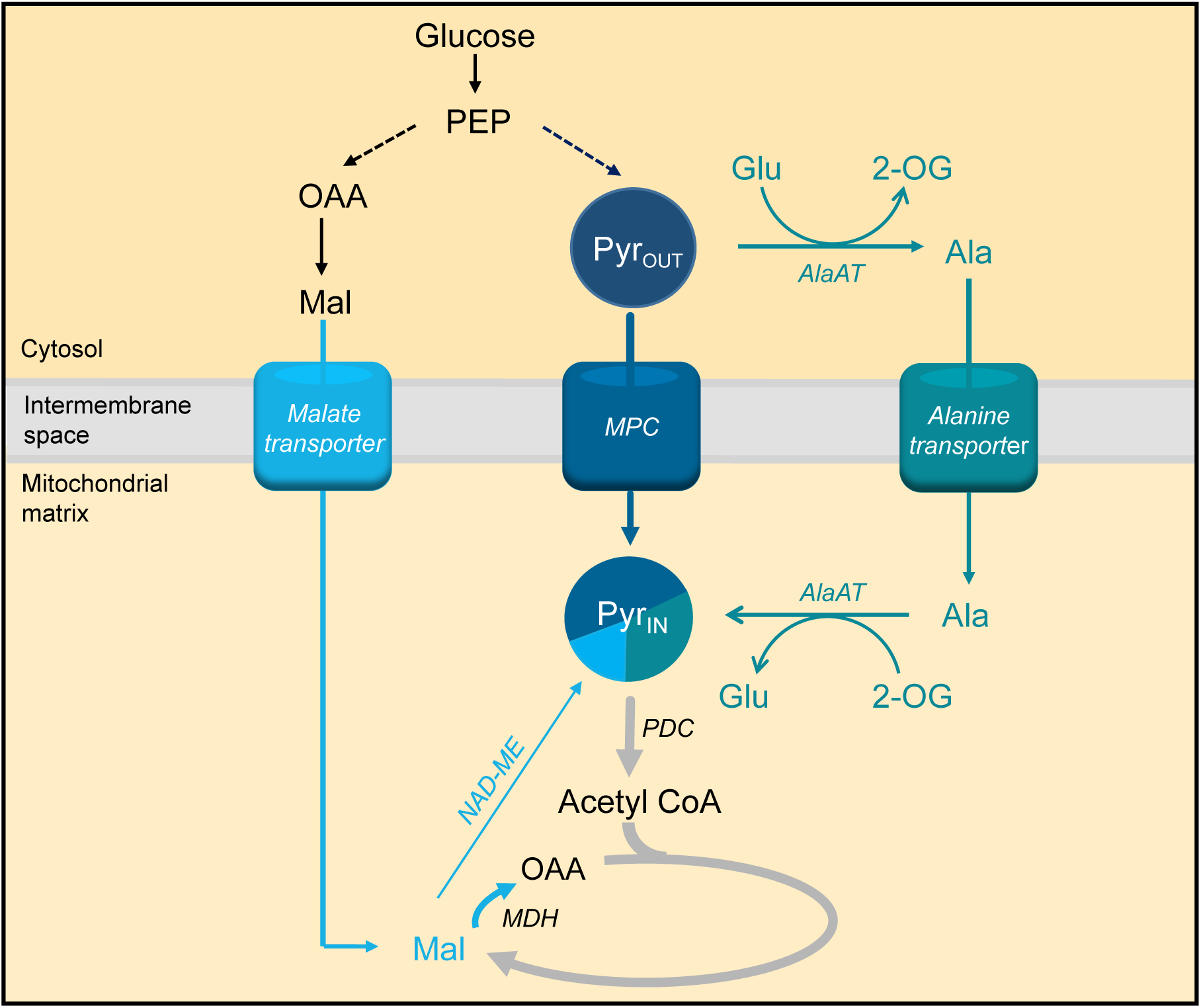
The mitochondrial pyruvate pool is contributed to by multiple supply pathways. A combination of *in vitro, in organello*, and *in vivo* analyses suggest the supply of mitochondrial pyruvate is facilitated mainly by MPC and AlaAT pathways; inhibition of one is compensated for by the other (e.g. increased alanine level is the most prominent feature of *mpc1* mutants). The majority of malate imported into the matrix under physiological pH conditions is likely to be oxidised into OAA, thus NAD-ME would synthesise a relatively small amount of pyruvate compared to the two major pathways, which was confirmed by analysis of *me1.me2*. MPC was the only pathway able to fully supply the pyruvate pool on its own, evidenced by the fact that cycloserine-treated *me1.me2* seedlings can maintain normal growth, while ALT alone in *me1.me2.mpc1* had a growth deficiency phenotype. Dashed lines indicate unknown relative fluxes. Bolder arrows indicate larger flux compared to other pathways using or generating the same metabolite. Abbreviations: AlaAT, alanine aminotransferase; MPC, mitochondrial pyruvate carrier; NAD-ME; NAD-malic enzyme; PDC, Pyruvate dehydrogenase complex; MDH, Malate dehydrogenase; OAA, oxaloacetate; PEP, Phosphoenolpyruvate; Pyr_out_, Pyruvate in the cytosol, Pyr _in_, Pyruvate inside the mitochondrial matrix; Ala, Alanine; Glu, Glutamate.

The degree to which MPC1 affects growth of different organisms reflects the flexibility of pyruvate metabolism and the proportion of oxidative phosphorylation generated by MPC-dependent pyruvate flux in different organisms. In mice, MPC is the primary carrier of oxidative substrates for energy production in mitochondria and catalyses most of the pyruvate flux, which cannot be sufficiently compensated for by other potential pyruvate-supplying pathways, leading to embryonic lethality when MPC is absent (Vanderperre et al., 2016). In yeast, the role of MPC is less important as the existence of a PDC bypass involving cytosolic pyruvate decarboxylation enables mitochondrial transport of substrates independently of MPC function (Boubekeur et al., 1999). Such flexibility of pyruvate utilisation also exists in Arabidopsis as seen by a lack of an obvious phenotype under normal growth conditions in mutant lacking the MPC complex (Supplementary Figure 1B). Our findings indicate that instead, plants utilise a mixture of MPC and MPC-bypasses to supply mitochondria with pyruvate. NAD-ME was more active in converting malate to pyruvate in isolated *mpc1* mitochondria compared to wildtype (Figure 2B and 3G). Conversely, knocking out NAD-ME resulted in an increase in MPC activity in isolated mitochondria as evidenced by the 50% increase in ^13^C incorporation into TCA intermediates (Figure 4E). The effect of impaired pyruvate supply on plant growth becomes visually apparent when MPC1 was knocked out in the *me1.me2* background. This triple mutant was not only unable to import pyruvate and convert malate to pyruvate but also showed a reduced the rate of mitochondrial malate transport, likely due to feedback inhibition by OAA accumulation in the matrix. Consistent with our results, knocking-out both MPC and mitochondrial NAD-ME in yeast magnified the slow-growth phenotype under non-stresses conditions which was not observed in the single MPC1 knock-out (Bricker et al., 2012). Taken together, the loss of MPC1 appears to trigger the promotion of NAD-ME flux to pyruvate and vice versa, at least in isolated mitochondria. This complementary regulation of MPC and mitochondrial NAD-ME activities is not as clear in single knockouts *in planta* due to metabolic flexibility which ensures that overall metabolic and energy homeostasis are maintained without plant growth consequences.

Metabolomics and ^13^C-pyruvate tracing in *vivo* only showed slight changes in metabolite levels in all genotypes examined, highlighting the metabolic flexibility of Arabidopsis leaves to maintain metabolite pool size in plants lacking MPC1. The only exceptions were alanine and 2-oxoglutarate abundances, which were higher in *mpc1* and *me1.me2* respectively. The phenotype of *me1.me2.mpc1* is most likely caused by the combined metabolic effects of these mutations, and the change in ^13^C labelling into glutamate (Figure 6B) which could be an indicator of an altered flux through AlaAT (Zhang et al., 2020). The hypersensitivity of the *me1.me2.mpc1* and *mpc1* plants to cycloserine treatment indicates that pyruvate–alanine cycling between mitochondria and cytosol can become a major pathway under certain conditions, such as when MPC1 is absent (Figure 7). However, there appears to be an energy cost associated with pyruvate-alanine cycling as the main mitochondrial pyruvate supply. The priority of diverting glutamate and 2-oxoglutarate pools into inter-conversion of pyruvate and alanine in mitochondria and cytosol makes these metabolites less available for other aminotransferases, which are critical for modulating cellular nitrogen metabolism and mediating nitrogen-dependent stress response (McCommis et al., 2015). Decreased cadmium sensitivity of *mpc1* (He et al., 2019) is an example of the consequences of increased flux through pyruvate-alanine cycling pathway as glutamate is not as readily available to produce glutathione to scavenge cadmium ions (Sytar et al., 2013; Tompkins et al., 2019) and to adjust amino acid profiles which have been shown to be important for heavy metal stress response (Blasco and Puppo, 1999; Sharma and Dietz, 2006; Zhu et al., 2018). Moreover, just as in MPC and NAD-ME single knockout mutants, mutant lacking the major AlaAT isoform (*alt1*) does not cause a vegetative phenotype (Miyashita et al., 2007), suggesting that at least two out of three pyruvate-supplying pathways are needed to support mitochondrial respiration and prevent an impact on plant growth.

Metabolic flexibility has long been considered a hallmark of what makes plant mitochondria unique (Douce and Neuburger, 1989). Much of the research on this flexibility has focused on the branching of the ETC (Vanlerberghe and McIntosh, 1997; Rasmusson et al., 2004; Millar et al., 2011) and the range of substrates that contribute reducing potential to the ETC (Schertl and Braun, 2014). Metabolic flexibility has been shown for TCA cycle components as well, given that these enzymes are mostly not exclusively targeted to mitochondria (Schnarrenberger and Martin, 2002; Araújo et al., 2012). Reduction in the activity of TCA cycle components such as mitochondrial citrate synthase (Sienkiewicz-Porzucek et al., 2008), NAD-dependent isocitrate dehydrogenase (Sienkiewicz-Porzucek et al., 2010), aconitase (Carrari et al., 2003), malate dehydrogenase (Nunes-Nesi et al., 2005) and fumarase (Nunes-Nesi et al., 2007) mostly resulted in normal phenotypes and minor changes in respiratory and photosynthetic performances in tomato plants due to metabolic re-adjustment through their counterparts in the cytosol, peroxisome and/or plastids. Furthermore, mitochondrial components often play an important role in a larger, highly flexible metabolic network linking carbon metabolism and nitrogen metabolism. For example, 2-oxoglutarate as the main checkpoint of both processes can be produced by the concerted action of isocitrate dehydrogenases, aminotransaminases, and glutamate dehydrogenase either in the mitochondrial matrix or cytosol (Weber and Flügge, 2002; Foyer et al., 2003). The appropriate allocation of 2-OG to TCA cycle and amino acid biosynthesis determines metabolic homeostasis. Also, the GABA shunt can bypass two steps of TCA cycle by allowing the synthesis of GABA in the cytosol during stresses including salinity, drought, heat, and hypoxia (Bouché and Fromm, 2004) and transport GABA back to the mitochondria to make succinate and re-enter the TCA cycle (Michaeli et al., 2011). In a similar manner, this work introduces AlaAT as a major flux contributor to the canonical pyruvate supply pathways by efficiently bypassing pyruvate import from the cytosol to the mitochondrial matrix rather than just being considered a stress response pathway (Ricoult et al., 2006; Miyashita and Good, 2008; Diab and Limami, 2016). In this context it is notable that a positive correlation of respiratory rates in Arabidopsis leaves was more apparent for alanine than for pyruvate, sugars or other organic and amino acids (O’Leary et al., 2017) and alanine is a strong stimulator of respiratory rate *in vivo* in Arabidopsis (O’Leary et al., 2020).

Mitochondrial pyruvate supply requires flexibility as pyruvate is an essential starting material for carbon metabolism in mitochondria, greatly affecting all downstream reactions and eventually energy production and plant development. The quantitative involvement of three pathways under changing internal and external environments in plants contributes to a remarkable plasticity in pyruvate supply in plants. While NAD-ME alone failed to compensate for the loss of both MPC and AlaAT, AlaAT alone (in *me1.me2.mpc1*) only resulted in a moderate vegetative phenotype while MPC alone (in cycloserine treated *me1.me2*) was able to maintain a normal plant growth (Figure 6). This suggests their relative contribution to the mitochondrial pyruvate pool *in vivo* and as a result their roles in central metabolism.

In conclusion, our data suggest plants possess at least three pathways that all synergistically contribute to mitochondrial pyruvate supply and usage for respiration. These pathways are coordinated in such a way that, when one is compromised the other two increase in flux to maintain metabolic and redox homeostasis for optimal growth. This allows metabolic flexibility to cope with changes in cellular substrate availability, particular during environmental changes such as nutrient and metal stresses, hypoxia, cold and drought. NAD-ME appears to contribute less to pyruvate supply than MPC and AlaAT pathways as AlaAT and MPC alone can largely support normal plant growth, but not NAD-ME alone (Figure 7). Meanwhile, AlaAT is an efficient alternative pathway to divert pyruvate flux from MPC especially when MPC is defective (Figure 7). A challenge for the future will be to accurately quantify the relative fluxes of all three pyruvate-contributing pathways to leaf respiration under different nutrient and stress conditions. This will be important to further understand one of the most important mitochondrial processes, and from there, the exploitation of plant energy generation and usage in plants.

## Methods

### Plant material, growth conditions and quantitative RNA analysis

The MPC1 T-DNA insertion line SALK008465 was obtained from the Arabidopsis Biological Resource Center (https://abrc.osu.edu/). *me1.me2* seeds were previously characterised and published (Tronconi et al., 2008) and were obtained from Professor Verónica G. Maurino (University of Bonn). *me1.me2.mpc1* was created from crossing between *me1.me2* and *mpc1. mpc1/gMPC1* and *me1.me2.mpc1/gMPC1* were generated by introducing the genomic version of *MPC1* with its endogenous promoter into the corresponding mutants using pCambia1380 as the vector. Specifically, genomic *MPC1* was amplified using gMPC1_F and gMPC1_R primers (Supplementary Table 1) and cloned into the plasmid pCambia1380 which was then transformed into *E. coli* to replicate the plasmid. The mutants were floral dipped in the culture of Agrobacterium transformed with pCambia1380-gMPC1. Seeds were collected and selected using 15 μg/mL hygromycin B plates for three generations before being utilized for experiments to ensure genetic stability.

Seeds were stratified for 3 days in the dark at 4°C before being transferred to either long day or short day conditions. Phenotype analysis was performed on plants grown under long day conditions of 16 hours light, 8 hours dark and 60% humidity (110 µmol s^−1^ m^−1^ light intensity). For cycloserine treatments, Arabidopsis seedlings were grown on vertical one-half strength Murashige and Skoog agar (10g/L agar, 0.4 g/L MES) plates with 0.5 µM cycloserine added after autoclaving.

Plants used in metabolic and stable isotope labelling analysis were grown under short day conditions of 8 hours light, 16 hours dark and 60% humidity (110 µmol s^−1^ m^−1^ light intensity).

RNA was extracted from three biological replicates per genotype using the Spectrum Plant Total RNA Kit (Sigma-Aldrich) according to the manufacturer’s instructions. The RNA quality and quantity were assessed using a nanodrop ND-1000 spectrophotometer (ThermoFisher). RNA was mixed with primers and qPCR reaction components provided in QuantiNova SYBR Green RT-PCR Kit in 96 well plate. Quantitative transcript analysis was done using the LightCycler® 480 System (Roche). A list of qPCR primers is provided in Supplementary Table 1.

### Isolation of mitochondria, O_2_ electrode measurements and enzyme activity assays

Arabidopsis seeds were surface sterilized and dispensed into one-half strength Murashige and Skoog liquid media (1.1g/L agar, 0.4 g/L MES, 10g/L sucrose) within enclosed, sterilized 100 mL polypropylene containers. The containers were seated on a rotating shaker in the long-day conditions as described above and seedlings were harvested after two weeks.

Mitochondria were isolated from 2-week old *Arabidopsis* seedlings as described previously (Millar et al., 2007). O_2_ electrode measurements were carried out as previously described (Lee et al., 2010), except pyruvate was added at 0, 0.5, 1, 2 or 5 mM and malate was added at 0.5 mM for pyruvate-dependent OCR measurement. PDH, NAD-ME *in vitro* activity assays were carried out as described previously (Huang et al., 2015).

### Analyses of metabolites by LC-SRM-MS

Leaf discs (∼20 mg) were collected from 6-weeks-old short-day grown (8-h light/16-h dark) plants at 1, 8, and 15 hours after dark switch and 4 hours after light switch. Samples were snap-frozen at −80°C and ground before metabolites were extracted using 90% methanol containing ^13^C-leucine and adipic acid as internal standards. After incubating at 75 °C and centrifugation at 18 000 *g* for three minutes, supernatants were aliquoted and dried using a SpeedVac concentrator.

For quantitation of organic acids, dried samples were resuspended in 100 μl methanol and derivatized as described previously (Han et al., 2013). Samples were analysed by an Agilent 1100 HPLC system coupled to an Agilent 6430 Triple Quadrupole (QQQ) mass spectrometer equipped with an electrospray ion source as described previously (Lee et al., 2020). Chromatographic separation was performed on a Kinetex C18 column, using 0.1% formic acid in water (solvent A) and 0.1% formic acid in methanol (solvent B) as the mobile phase for binary gradient elution (Supplementary Table 2). Data acquisition was performed using Agilent MassHunter Workstation Data Acquisition software. The column flow rate was 0.3 mL/min; the column temperature is 40 °C, and the autosampler was kept at 10 °C. Selective reaction monitoring (SRM) transitions for each targeted analyte are shown in Supplementary Table 3. Optimisations for metabolite transitions for SRM-MS were described in detail elsewhere (Le et al., 2021).

For amino acid quantification, dried samples were resuspended in 50 μl water. Samples were analysed using the same LC-MS system as described above. Chromatographic separation was performed using Agilent Poroshell 120 HILIC-Z column, using mobile phases of 20mM ammonium formate in water (solvent A) and 20 mM ammonium formate in acetonitrile (Solvent B, Supplementary Table 4). The column flow rate was 0.4 mL/min; the column temperature was 35 °C, and the autosampler was kept at 10°C. SRM transitions for each targeted analyte are listed in Supplementary Table 5.

### Substrate feeding of isolated mitochondria

The detailed methods and materials for MS-based mitochondria feeding assays were described previously (Le et al., 2021). In short, 100 µg isolated mitochondria were mixed with substrates (0.5mM pyruvate and 0.5mM malate), cofactors (2 mM NAD^+^, 0.2 mM TPP and 0.012 mM CoA) and 1 mM ADP (for ATP synthesis) in a final volume of 200 µl. At specified times, this reaction mixture was layered on top of silicon oil (AR200, 90 µl) which was layered above the stopping sucrose solution (0.5 M sucrose, pH 1.0). Substrate transport was stopped by rapid centrifugation (12 000 *g* for 3 min) to harvest the mitochondria at the bottom of the tube. 5 µl of the extra-mitochondrial medium (the top layer) was collected and extracted for quantitative analysis by LC-SRM-MS.

### ^13^C-pyruvate and ^13^C-glucose labelling of Arabidopsis leaf discs and analysis of labelled metabolites

Leaf discs (∼50 mg) were prepared from 6-weeks-old short-day grown (8-h light/16-h dark) plants 1 hour before the end of a normal light photoperiod. They were floated on leaf respiratory buffer containing 20 mM ^13^C_3_-pyruvate (99% purity, Sigma Aldrich), 20 mM ^13^C_6_-glucose (99% purity, Sigma Aldrich) or an unlabelled counterpart. At the specified incubation time, leaf discs were briefly washed with respiratory buffer to remove excess labelled pyruvate and frozen in liquid nitrogen for metabolite extraction as stated above. Analyses of total, untargeted metabolites were performed using an Agilent 1100 HPLC system coupled to an Agilent 6520 Quadrupole/Time-of-Flight (Q-TOF) mass spectrometer equipped with an electrospray ion source. Chromatographic separation was performed on an Agilent Poroshell 120 HILIC-Z column, using 10 mM ammonium acetate in water (solvent A) and 10 mM ammonium acetate in acetonitrile (Solvent B) as mobile phases (Supplementary Table 6). Solvent A and B were supplied with 0.1% (v/v) Infinity Lab deactivator additive (Agilent) to improve peak shapes. The column flow rate was 0.25 mL/min; the column temperature was 40 °C, and the autosampler was kept at 10 °C. The scanning range was from 70 m/z to 230 m/z.

Peak extraction was done using Agilent Mass Hunter Quantitative analysis software based on known m/z values and retention time obtained from authentic standards. Raw peak was then isotopically corrected for natural ^12^C abundances using IsoCor software (Millard et al., 2019). Percentage of each isotopologue of a compound of interest was calculated as the ratio of the peak area of the isotopologue over the total area of both labelled and unlabelled fractions. Absolute labelled amount of each isotopologue of a compound was calculated through multiplying the percentage of labelled isotopologue by the absolute amount of that compound obtained from the analysis of unlabelled sample by LC-SRM-MS as described above.

### Relative quantification of mitochondrial protein abundances by LC-SRM-MS

100 µg frozen mitochondrial proteins were precipitated in 100% acetone for 24 hours at - 20°C and the pellets were washed with acetone for three times. Samples were alkylated, trypsin digested, desalted and cleaned as previously described (Petereit et al., 2020). Samples were loaded onto an AdvanceBio Peptide Map column (2.1 × 250 mm, 2.7 μm particle size; part number 651750-902, Agilent), using an Agilent 1290 Infinity II LC System coupled to an Agilent 6495 Triple Quadrupole MS. The column was heated to 60°C and the column flow rate was 0.4 mL/min. The binary elution gradient for HPLC and the list of peptides transitions used for SRM-MS are provided in Supplementary Table 7 and 8 respectively. Peak area of targeted peptides were determined using the Skyline software package. Peptide abundances from each sample were normalised against VDAC since its abundance was similar amongst all samples.

### Statistical Analysis

All statistical analyses were performed using the T.Test function built in Excel 2010. Statistical tests and the number of biological replicates are indicated in figure legends. Biological replicates indicate samples that were collected from different batches plants grown under the same conditions except biological replicates for transcript analysis and metabolite analysis were samples collected from different plants grown at the same time.

### Accession Numbers

Sequence data from this article can be found in the Arabidopsis Genome Initiative or GenBank/EMBL databases under the following accession numbers: AtMPC1 (At5g20090), AtMPC4 (At4g05590), AtMPC3 (At4g22310), AtMPC2 (At4g14695), AtNAD-ME1 (At2g13560) and AtNAD-ME2 (At4g00570).

## Supplemental Data

**Supplemental Figure 1:** Gene model and phenotypes of *mpc1*

**Supplemental Figure 2:** Changes in the abundance and activity of respiratory enzymes in tested mutants

**Supplemental Figure 3:** ^13^C_3_-Pyruvate and malate feeding to isolated mitochondria of Col-0, *mpc1* and *mpc1/gMPC1*

**Supplemental Figure 4:** Swap label experiment of pyruvate and ^13^C_4_-malate feeding into isolated mitochondria of Col-0, *mpc1* and *mpc1/gMPC1*

**Supplemental Figure 5:** Calculated import and export rates from the swap label time course experiment of pyruvate and ^13^C_4_-malate feeding into isolated mitochondria of Col-0, *mpc1* and *mpc1/gMPC1*

**Supplemental Figure 6:** Soil-based phenotypes of *mpc1*, *me1.me2* and complementary genotypes

**Supplemental Figure 7:** Time courses of metabolite concentrations in the extra-mitochondrial space of ^13^C_3_-Pyruvate and malate feeding into *Col-0, me1.me2* and *me1.me2.mpc1* mitochondria

**Supplementary Figure 8:** Diurnal changes of representative metabolites in different genotypes

**Supplementary Figure 9:** Incorporation of ^13^C_3_-pyruvate and ^13^C-glucose into alanine, TCA cycle intermediates and related amino acids of leaf discs from different genotypes in the dark.

**Supplementary Table 1:** A list of qPCR primers used.

**Supplementary Table 2:** The binary elution gradient of solvent A and solvent B for HPLC used for organic acid quantification.

**Supplementary Table 3:** The SRM transitions used as quantifier and qualifier for derivatised labelled and unlabelled TCA cycle metabolites of interest.

**Supplementary Table 4:** The binary elution gradient of solvent A and solvent B for HPLC used for amino acid quantification.

**Supplementary Table 5:** The SRM transitions used as quantifier and qualifier for amino acid measurement.

**Supplementary Table 6:** The binary elution gradient of solvent A and solvent B for HPLC used for untargeted metabolite analysis.

**Supplementary Table 7:** The binary elution gradient of solvent A and solvent B for HPLC used for mitochondrial protein abundance measurement.

**Supplementary Table 8:** The SRM transitions used to measure relative abundance of mitochondrial proteins.

## Acknowledgements

We thank Ricarda Fenske for optimising method and running LC-SRM-MS for the mitochondrial protein abundance measurements. We thank Brendan O’Leary for a critical reading of the manuscript. This work is supported by the Australian Research Council Centre of Excellence in Plant Energy Biology (CE140100008) and X.L. is a Forrest Scholar supported by the Forrest Research Foundation and a receiver of Research Training Program scholarships from the Department of Education, Skills and Employment in the Australian Government.

## Author contributions

XL, CPL and AHM designed the research. XL performed most of the experiments and data analysis, CPL assisted with some of the mass spectrometry and data analysis. XL, CPL and AHM wrote the paper.

## REFERENCES

Araújo, W.L., Nunes-Nesi, A., Nikoloski, Z., Sweetlove, L.J., And Fernie, A.R. (2012). Metabolic control and regulation of the tricarboxylic acid cycle in photosynthetic and heterotrophic plant tissues. Plant Cell Environ 35, 1–21.

Brailsford, M.A., Thompson, A.G., Kaderbhai, N., and Beechey, R.B. (1986). Pyruvate metabolism in castor-bean mitochondria. Biochem J 239, 355–361.

Bender, T., and Martinou, J.C. (2016). The mitochondrial pyruvate carrier in health and disease: To carry or not to carry? Biochim Biophys Acta 1863, 2436–2442.

Blasco, J., and Puppo, J. (1999). Effect of heavy metals (Cu, Cd and Pb) on aspartate and alanine aminotransferase in *Ruditapes philippinarum* (Mollusca: Bivalvia). Comp Biochem Physiol C Pharmacol Toxicol Endocrinol 122, 253–263.

Boubekeur, S., Bunoust, O., Camougrand, N., Castroviejo, M., Rigoulet, M., and Guérin, B. (1999). A mitochondrial pyruvate dehydrogenase bypass in the yeast *Saccharomyces cerevisiae*. J Biol Chem 274, 21044–21048.

Bouché, N., and Fromm, H. (2004). GABA in plants: just a metabolite? Trends Plant Sci 9, 110–115.

Bowsher, C.G., and Tobin, A.K. (2001). Compartmentation of metabolism within mitochondria and plastids. J Exp Bot 52, 513–527.

Bricker, D.K., Taylor, E.B., Schell, J.C., Orsak, T., Boutron, A., Chen, Y.C., Cox, J.E., Cardon, C.M., Van Vranken, J.G., Dephoure, N., Redin, C., Boudina, S., Gygi, S.P., Brivet, M., Thummel, C.S., and Rutter, J. (2012). A mitochondrial pyruvate carrier required for pyruvate uptake in yeast, Drosophila, and humans. Science 337, 96.

Brivet, M., Garcia-Cazorla, A., Lyonnet, S., Dumez, Y., Nassogne, M.C., Slama, A., Boutron, A., Touati, G., Legrand, A., and Saudubray, J.M. (2003). Impaired mitochondrial pyruvate importation in a patient and a fetus at risk. Mol Genet Metab 78, 186–192.

Carrari, F., Nunes-Nesi, A., Gibon, Y., Lytovchenko, A., Loureiro, M.E., and Fernie, A.R. (2003). Reduced expression of aconitase results in an enhanced rate of photosynthesis and marked shifts in carbon partitioning in illuminated leaves of wild species tomato. Plant Physiol 133, 1322–1335.

Chinopoulos, C. (2020). From Glucose to Lactate and Transiting Intermediates Through Mitochondria, Bypassing Pyruvate Kinase: Considerations for Cells Exhibiting Dimeric PKM2 or Otherwise Inhibited Kinase Activity. Front Physiol 11.

Cornell, N.W., Zuurendonk, P.F., Kerich, M.J., and Straight, C.B. (1984). Selective inhibition of alanine aminotransferase and aspartate aminotransferase in rat hepatocytes. Biochem J 220, 707–716.

Day, D.A. (1980). Malate Decarboxylation by *Kalanchoë daigremontiana* Mitochondria and Its Role in Crassulacean Acid Metabolism. Plant Physiol 65, 675–679.

Day, D.A., and Hanson, J.B. (1977). Pyruvate and malate transport and oxidation in corn mitochondria. Plant Physiol 59, 630–635.

Diab, H., and Limami, A.M. (2016). Reconfiguration of N metabolism upon hypoxia stress and recovery: Roles of alanine aminotransferase (AlaAT) and glutamate dehydrogenase (GDH). Plants (Basel) 5, 25.

Douce, R., and Neuburger, M. (1989). The uniqueness of plant mitochondria. Annu Rev of Plant Physiol Plant Mol Biol 40, 371–414.

Duff, S.M.G., Rydel, T.J., McClerren, A.L., Zhang, W., Li, J.Y., Sturman, E.J., Halls, C., Chen, S., Zeng, J., Peng, J., Kretzler, C.N., and Evdokimov, A. (2012). The enzymology of alanine aminotransferase (AlaAT) isoforms from *Hordeum vulgare* and other organisms, and the HvAlaAT crystal structure. Arch of Biochem and Biophys 528, 90–101.

Edwards, S., Nguyen, B.T., Do, B., and Roberts, J.K.M. (1998) Contribution of malic enzyme, pyruvate kinase, phosphoenolpyruvate carboxylase, and the krebs cycle to respiration and biosynthesis and to intracellular pH regulation during hypoxia in maize root tips observed by nuclear magnetic resonance imaging and gas chromatography-mass spectrometry. Plant Physiol 116, 1073–1081.

Foyer, C.H., Parry, M., and Noctor, G. (2003). Markers and signals associated with nitrogen assimilation in higher plants. J Exp Bot 54, 585–593.

Haferkamp, I., and Schmitz-Esser, S. (2012). The Plant Mitochondrial Carrier Family: Functional and Evolutionary Aspects. Front in Plant Sci 3.

Halestrap, A.P. (1975). The mitochondrial pyruvate carrier. Kinetics and specificity for substrates and inhibitors. Biochem J 148, 85–96.

Han, J., Gagnon, S., Eckle, T., and Borchers, C.H. (2013). Metabolomic analysis of key central carbon metabolism carboxylic acids as their 3-nitrophenylhydrazones by UPLC/ESI-MS. Electrophoresis 34, 2891–2900.

Hartwell, J., Dever, L.V., and Boxall, S.F. (2016). Emerging model systems for functional genomics analysis of Crassulacean acid metabolism. Curr Opin Plant Biol 31, 100–108.

He, L., Jing, Y., Shen, J., Li, X., Liu, H., Geng, Z., Wang, M., Li, Y., Chen, D., Gao, J., and Zhang, W. (2019). Mitochondrial pyruvate carriers prevent cadmium toxicity by sustaining the TCA cycle and glutathione synthesis. Plant Physiol 180, 198.

Herzig, S., Raemy, E., Montessuit, S., Veuthey, J.-L., Zamboni, N., Westermann, B., Kunji, E.R.S., and Martinou, J.C. (2012). Identification and functional expression of the mitochondrial pyruvate carrier. Science 337, 93.

Hong, C.S., Graham, N.A., Gu, W., Espindola Camacho, C., Mah, V., Maresh, E.L., Alavi, M., Bagryanova, L., Krotee, P.A.L., Gardner, B.K., Behbahan, I.S., Horvath, S., Chia, D., Mellinghoff, I.K., Hurvitz, S.A., Dubinett, S.M., Critchlow, S.E., Kurdistani, S.K., Goodglick, L., Braas, D., Graeber, T.G., and Christofk, H.R. (2016). MCT1 Modulates Cancer Cell Pyruvate Export and Growth of Tumors that Co-express MCT1 and MCT4. Cell Rep 14, 1590–1601.

Huang, S., Lee, C.P., and Millar, A.H. (2015). Activity assay for plant mitochondrial enzymes. Methods Mol Biol 1305, 139–149.

Hüdig, M., Maier, A., Scherrers, I., Seidel, L., Jansen, E.E.W., Mettler-Altmann, T., Engqvist, M.K.M., and Maurino, V.G. (2015). Plants Possess a Cyclic Mitochondrial Metabolic Pathway similar to the Mammalian Metabolic Repair Mechanism Involving Malate Dehydrogenase and l-2-Hydroxyglutarate Dehydrogenase. Plant Cell Physiol 56, 1820–1830.

Jenner, H.L., Winning, B.M., Millar, A.H., Tomlinson, K.L., Leaver, C.J., and Hill, S.A. (2001). NAD malic enzyme and the control of carbohydrate metabolism in potato tubers. Plant Physiol 126, 1139–1149.

Le, X., Millar, A.H., and Lee, C.P. (2021). Assessing the kinetics of metabolite uptake and utilisation by isolated mitochondria using selective reaction monitoring-mass spectrometry (SRM-MS). In Methods in Molecular Biology: Plant Mitochondria (Springer), accepted

Lee, C.P., Eubel, H., and Millar, A.H. (2010). Diurnal changes in mitochondrial function reveal daily optimization of light and dark respiratory metabolism in Arabidopsis. Mol Cell Proteomics 9, 2125–2139.

Lee, C.P., Elsässer, M., Fuchs, P., Fenske, R., Schwarzländer, M., and Millar, A.H. (2020). The Arabidopsis mitochondrial dicarboxylate carrier 2 maintains leaf metabolic homeostasis by uniting malate import and citrate export. bioRxiv, https://doi.org/10.1101/2020.04.28.065441.

Lehmann, M.M., Wegener, F., Barthel, M., Maurino, V.G., Siegwolf, R.T.W., Buchmann, N., Werner, C., and Werner, R.A. (2016). Metabolic Fate of the Carboxyl Groups of Malate and Pyruvate and their Influence on δ13C of Leaf-Respired CO_2_ during Light Enhanced Dark Respiration. Front Plant Sci 7.

Li, C.L., Wang, M., Ma, X.Y., and Zhang, W. (2014). NRGA1, a putative mitochondrial pyruvate carrier, mediates ABA regulation of guard cell ion channels and drought stress responses in Arabidopsis. Mol Plant 7, 1508–1521.

Macrae, A.R., and Moorhouse, R. (1970). The oxidation of malate by mitochondria isolated from cauliflower buds. Eur J Biochem 16, 96–102.

McCommis, K.S., Chen, Z., Fu, X., McDonald, W.G., Colca, J.R., Kletzien, R.F., Burgess, S.C., and Finck, B.N. (2015). Loss of mitochondrial pyruvate carrier 2 in the liver leads to defects in gluconeogenesis and compensation via pyruvate-alanine cycling. Cell Metab 22, 682–694.

Michaeli, S., Fait, A., Lagor, K., Nunes-Nesi, A., Grillich, N., Yellin, A., Bar, D., Khan, M., Fernie, A.R., Turano, F.J., and Fromm, H. (2011). A mitochondrial GABA permease connects the GABA shunt and the TCA cycle, and is essential for normal carbon metabolism. Plant J 67, 485–498.

Millar, A.H., Liddell, A., and Leaver, C.J. (2007). Isolation and Subfractionation of Mitochondria from Plants. In Methods in Cell Biology (Academic Press), pp. 65–90.

Millar, A.H., Whelan, J., Soole, K.L., and Day, D.A. (2011). Organization and regulation of mitochondrial respiration in plants. Annu Rev Plant Biol 62, 79–104.

Millard, P., Delépine, B., Guionnet, M., Heuillet, M., Bellvert, F., and Létisse, F. (2019). IsoCor: isotope correction for high-resolution MS labeling experiments. Bioinformatics 35, 4484–4487.

Miyashita, Y., and Good, A.G. (2008). Contribution of the GABA shunt to hypoxia-induced alanine accumulation in roots of *Arabidopsis thaliana*. Plant Cell Physiol 49, 92–102.

Miyashita, Y., Dolferus, R., Ismond, K.P., and Good, A.G. (2007). Alanine aminotransferase catalyses the breakdown of alanine after hypoxia in Arabidopsis thaliana. Plant J 49, 1108–1121.

Netting, A.G. (2002). pH, abscisic acid and the integration of metabolism in plants under stressed and non-stressed conditions. II. Modifications in modes of metabolism induced by variation in the tension on the water column and by stress. J Exp Bot 53, 151–173.

Neuburger, M., and Douce, R. (1980). Effect of bicarbonate and oxaloacetate on malate oxidation by spinach leaf mitochondria. Biochim et Biophys Acta 589, 176–189.

Nunes-Nesi, A., Carrari, F., Lytovchenko, A., Smith, A.M.O., Ehlers Loureiro, M., Ratcliffe, R.G., Sweetlove, L.J., and Fernie, A.R. (2005). Enhanced photosynthetic performance and growth as a consequence of decreasing mitochondrial malate dehydrogenase activity in transgenic tomato plants. Plant Physiol 137, 611–622.

Nunes-Nesi, A., Carrari, F., Gibon, Y., Sulpice, R., Lytovchenko, A., Fisahn, J., Graham, J., Ratcliffe, R.G., Sweetlove, L.J., and Fernie, A.R. (2007). Deficiency of mitochondrial fumarase activity in tomato plants impairs photosynthesis via an effect on stomatal function. Plant J 50, 1093–1106.

O’Leary, B.M., Oh, G.G.K., Lee, C.P., and Millar, A.H. (2020). Metabolite regulatory interactions control plant respiratory metabolism via target of rapamycin (TOR) kinase activation. Plant Cell 32, 666–682.

O’Leary, B.M., Lee, C.P., Atkin, O.K., Cheng, R., Brown, T.B., and Millar, A.H. (2017). Variation in leaf respiration rates at night correlates with carbohydrate and amino acid supply. Plant Physiol 174, 2261–2273.

Ohbayashi, I., Huang, S., Fukaki, H., Song, X., Sun, S., Morita, M.T., Tasaka, M., Millar, A.H., and Furutani, M. (2019). Mitochondrial pyruvate dehydrogenase contributes to auxin-regulated organ development. Plant Physiol 180, 896.

Passarella, S., Atlante, A., Valenti, D., and de Bari, L. (2003). The role of mitochondrial transport in energy metabolism. Mitochondrion 2, 319–343.

Petereit, J., Duncan, O., Murcha, M.W., Fenske, R., Cincu, E., Cahn, J., Pružinská, A., Ivanova, A., Kollipara, L., Wortelkamp, S., Sickmann, A., Lee, J., Lister, R., Millar, A.H., and Huang, S. (2020). Mitochondrial CLPP2 assists coordination and homeostasis of respiratory complexes. Plant Physiol 184, 148–164.

Rao, X., and Dixon, R.A. (2016). The Differences between NAD-ME and NADP-ME Subtypes of C_4_ Photosynthesis: More than Decarboxylating Enzymes. Front Plant Sci 7, 1525–1525.

Rasmusson, A.G., Soole, K.L., and Elthon, T.E. (2004). Alternative NAD(P)H dehydrogenases of plant mitochondria. Annu Rev Plant Biol 55, 23–39.

Ricoult, C., Echeverria, L.O., Cliquet, J.B., and Limami, A.M. (2006). Characterization of alanine aminotransferase (AlaAT) multigene family and hypoxic response in young seedlings of the model legume *Medicago truncatula*. J Exp Bot 57, 3079–3089.

Schertl, P., and Braun, H.-P. (2014). Respiratory electron transfer pathways in plant mitochondria. Front Plant Sci 5, 163.

Schmidtmann, E., König, A.-C., Orwat, A., Leister, D., Hartl, M., and Finkemeier, I. (2014). Redox regulation of Arabidopsis mitochondrial citrate synthase. Mol Plant 7, 156–169.

Schnarrenberger, C., and Martin, W. (2002). Evolution of the enzymes of the citric acid cycle and the glyoxylate cycle of higher plants. Eur J Biochem 269, 868–883.

Schwacke, R., Schneider, A., van der Graaff, E., Fischer, K., Catoni, E., Desimone, M., Frommer, W.B., Flügge, U.-I., and Kunze, R. (2003). ARAMEMNON, a novel database for Arabidopsis integral membrane proteins. Plant Physiol 131, 16–26.

Selinski, J., and Scheibe, R. (2019). Malate valves: old shuttles with new perspectives. Plant Biol 21, 21–30.

Sharma, S.S., and Dietz, K.J. (2006). The significance of amino acids and amino acid-derived molecules in plant responses and adaptation to heavy metal stress. J Exp Bot 57, 711–726.

Shen, J.L., Li, C.L., Wang, M., He, L.L., Lin, M.Y., Chen, D.H., and Zhang, W. (2017). Mitochondrial pyruvate carrier 1 mediates abscisic acid-regulated stomatal closure and the drought response by affecting cellular pyruvate content in *Arabidopsis thaliana*. BMC Plant Biol 17, 217.

Sienkiewicz-Porzucek, A., Nunes-Nesi, A., Sulpice, R., Lisec, J., Centeno, D.C., Carillo, P., Leisse, A., Urbanczyk-Wochniak, E., and Fernie, A.R. (2008). Mild reductions in mitochondrial citrate synthase activity result in a compromised nitrate assimilation and reduced leaf pigmentation but have no effect on photosynthetic performance or growth. Plant Physiol 147, 115–127.

Sienkiewicz-Porzucek, A., Sulpice, R., Osorio, S., Krahnert, I., Leisse, A., Urbanczyk-Wochniak, E., Hodges, M., Fernie, A.R., and Nunes-Nesi, A. (2010). Mild reductions in mitochondrial nad-dependent isocitrate dehydrogenase activity result in altered nitrate assimilation and pigmentation but do not impact growth. Mol Plant 3, 156–173.

Singh, B.K., and Shaner, D.L. (1995). Biosynthesis of branched chain amino acids: from test tube to field. Plant Cell 7, 935–944.

Sytar, O., Kumar, A., Latowski, D., Kuczynska, P., Strzałka, K., and Prasad, M.N.V. (2013). Heavy metal-induced oxidative damage, defense reactions, and detoxification mechanisms in plants. Acta Physiole Plant 35, 985–999.

Tavoulari, S., Thangaratnarajah, C., Mavridou, V., Harbour, M.E., Martinou, J.C., and Kunji, E.R. (2019). The yeast mitochondrial pyruvate carrier is a hetero-dimer in its functional state. EMBO J 38.

Tcherkez, G., Cornic, G., Bligny, R., Gout, E., and Ghashghaie, J. (2005). *In vivo* respiratory metabolism of illuminated leaves. Plant Physiol 138, 1596–1606.

Tcherkez, G., Bligny, R., Gout, E., Mahé, A., Hodges, M., and Cornic, G. (2008). Respiratory metabolism of illuminated leaves depends on CO_2_ and O_2_ conditions. Proc Natl Acad Sci 105, 797–802.

Tcherkez, G., Mahé, A., Gauthier, P., Mauve, C., Gout, E., Bligny, R., Cornic, G., and Hodges, M. (2009). In Folio Respiratory Fluxomics Revealed by ^13^C Isotopic Labeling and H/D Isotope Effects Highlight the Noncyclic Nature of the Tricarboxylic Acid “Cycle” in Illuminated Leaves. Plant Physiol 151, 620–630.

Tomaz, T., Bagard, M., Pracharoenwattana, I., Lindén, P., Lee, C.P., Carroll, A.J., Ströher, E., Smith, S.M., Gardeström, P., and Millar, A.H. (2010). Mitochondrial malate dehydrogenase lowers leaf respiration and alters photorespiration and plant growth in Arabidopsis. Plant Physiol 154, 1143–1157.

Tompkins, S.C., Sheldon, R.D., Rauckhorst, A.J., Noterman, M.F., Solst, S.R., Buchanan, J.L., Mapuskar, K.A., Pewa, A.D., Gray, L.R., Oonthonpan, L., Sharma, A., Scerbo, D.A., Dupuy, A.J., Spitz, D.R., and Taylor, E.B. (2019). Disrupting mitochondrial pyruvate uptake directs glutamine into the TCA cycle away from glutathione synthesis and impairs hepatocellular tumorigenesis. Cell Rep 28, 2608–2619.e2606.

Tronconi, M.A., Maurino, V.G., Andreo, C.S., and Drincovich, M.F. (2010a). Three different and tissue-specific NAD-malic enzymes generated by alternative subunit association in *Arabidopsis thaliana*. J Biol Chem 285, 11870–11879.

Tronconi, Marcos A., Gerrard Wheeler, Mariel C., Maurino, Verónica G., Drincovich, María F., and Andreo, Carlos S. (2010b). NAD-malic enzymes of *Arabidopsis thaliana* display distinct kinetic mechanisms that support differences in physiological control. Biochem J 430, 295–303.

Tronconi, M.A., Fahnenstich, H., Gerrard Weehler, M.C., Andreo, C.S., Flügge, U.I., Drincovich, M.F., and Maurino, V.G. (2008). Arabidopsis NAD-malic enzyme functions as a homodimer and heterodimer and has a major impact on nocturnal metabolism. Plant Physiol 146, 1540–1552.

Vanderperre, B., Herzig, S., Krznar, P., Hörl, M., Ammar, Z., Montessuit, S., Pierredon, S., Zamboni, N., and Martinou, J.C. (2016). Embryonic lethality of mitochondrial pyruvate carrier 1 deficient mouse can be rescued by a ketogenic diet. PLoS Genet 12, e1006056.

Vanlerberghe, G.C., and McIntosh, L. (1997). Alternative oxidase: From Gene to Function. Annu Rev Plant Physiol Plant Mol Biol 48, 703–734.

Waese, J., Fan, J., Pasha, A., Yu, H., Fucile, G., Shi, R., Cumming, M., Kelley, L.A., Sternberg, M.J., Krishnakumar, V., Ferlanti, E., Miller, J., Town, C., Stuerzlinger, W., and Provart, N.J. (2017). ePlant: Visualizing and exploring multiple levels of data for hypothesis generation in plant biology. Plant Cell 29, 1806–1821.

Weber, A., and Flügge, U.I. (2002). Interaction of cytosolic and plastidic nitrogen metabolism in plants. J Exp Bot 53, 865–874.

Willeford, K., and Wedding, R. (1987). pH Effects on the Activity and Regulation of the NAD malic enzyme. Plant Physiol 84, 1084–1087.

Wong, D.T., Fuller, R.W., and Molloy, B.B. (1973). Inhibition of amino acid transaminases by L-cycloserine. Adv Enzyme Regul 11, 139–154.

Zhang, Y., Taufalele, P.V., Cochran, J.D., Robillard-Frayne, I., Marx, J.M., Soto, J., Rauckhorst, A.J., Tayyari, F., Pewa, A.D., Gray, L.R., Teesch, L.M., Puchalska, P., Funari, T.R., McGlauflin, R., Zimmerman, K., Kutschke, W.J., Cassier, T., Hitchcock, S., Lin, K., Kato, K.M., Stueve, J.L., Haff, L., Weiss, R.M., Cox, J.E., Rutter, J., Taylor, E.B., Crawford, P.A., Lewandowski, E.D., Des Rosiers, C., and Abel, E.D. (2020). Mitochondrial pyruvate carriers are required for myocardial stress adaptation. Nat Metab 2, 1248–1264.

Zhu, G., Xiao, H., Guo, Q., Zhang, Z., Zhao, J., and Yang, D. (2018). Effects of cadmium stress on growth and amino acid metabolism in two Compositae plants. Ecotoxicol Environ Saf 158, 300–308.

